# Evaluation of normalization methods for predicting quantitative phenotypes in metagenomic data analysis

**DOI:** 10.1101/2023.10.25.563937

**Authors:** Beibei Wang, Yihui Luan

**Affiliations:** Frontier Science Center for Nonlinear Expectations, Ministry of Education, Qingdao, Shandong, China; Research Center for Mathematics and Interdisciplinary Sciences, Shandong University, Qingdao, Shandong, China; School of Mathematics, Shandong University, Jinan, Shandong, China

## Abstract

Significant advancements have been made in metagenomic research for the prediction of phenotypes based on microbiome data. While qualitative case-control predictions have received significant attention, less emphasis has been placed on predicting quantitative phenotypes. This emerging field holds great promise in revealing intricate connections between microbial communities and host health. However, the presence of heterogeneity in microbiome datasets poses a substantial challenge to the accuracy of predictions and undermines the reproducibility of models. To tackle this challenge, we investigated 22 normalization methods aimed at removing heterogeneity across multiple datasets, conducted a comprehensive review of them, and evaluated their effectiveness in predicting quantitative phenotypes using 3 simulations and 31 real datasets. The results indicate that none of these methods demonstrate significant superiority in predicting quantitative phenotypes or attain a noteworthy reduction in root mean squared error of the predictions. Given the frequent occurrence of batch effects and the satisfactory performance of batch correction methods in predicting datasets affected by these effects, we strongly recommend utilizing batch correction methods as the initial step in predicting quantitative phenotypes. In summary, the performance of normalization methods in predicting metagenomic data remains a dynamic and ongoing research area. Our study contributes to this field by undertaking a comprehensive evaluation of diverse methods and offering valuable insights into their effectiveness in predicting quantitative phenotypes.

## Introduction

The microbiome, which encompasses a complex ecosystem of microorganisms inhabiting the human body, plays a vital role in both host health and disease [1]. Understanding the intricate interplay between microbial communities and host phenotypes has substantial implications for enhancing our knowledge of diverse health-related conditions.

The utilization of microbiome data to predict phenotypes has become increasingly important in the era of high-throughput sequencing and metagenomics. A considerable amount of research in metagenomics has traditionally focused on predicting qualitative phenotypes, such as disease and non-disease states [2, 3]. However, there has been comparatively less emphasis on predicting quantitative phenotypes, which include numerical and continuous traits such as Body Mass Index (BMI) or blood glucose levels. The prediction framework for quantitative phenotypes is currently receiving increasing attention due to its significance. For instance, Yun et al. [4] identified distinct differences in gut microbiome composition among individuals with varying BMIs, providing valuable insights into the influence of microbial communities on body weight. In another study, Krisko et al. [5] suggested that the gut microbiome plays a role in regulating blood glucose levels, presenting opportunities for personalized interventions and treatments. Therefore, exploring the associations between the microbiome and quantitative health-related phenotypes is essential for unraveling the intricate interplay between the microbiome and human health, an area that has not been well addressed.

Quantitative phenotype prediction using microbiome data has its challenges. One of the foremost obstacles is the heterogeneity present in different datasets. Variations in geographic origin, sampling protocols, sequencing platforms, and other technical factors contribute to this heterogeneity, diminishing the generalizability and predictive accuracy of models. To enhance the reproducibility of predictive models, many studies have been dedicated to mitigating heterogeneity in predictions. These studies often involve merging data from distinct datasets into one and treating them as if they originate from the same dataset to improve prediction accuracy [3, 6]. Alternatively, researchers integrate the trained predictors from different datasets using diverse strategies to generate enhanced predictions [7, 8]. However, these techniques face limitations when dealing with datasets of limited sample size. In such a process, the potential contribution of normalization methods in prediction has often been overlooked.

Normalization serves as a crucial preprocessing step, effectively reducing data heterogeneity and enhancing the comparability of microbiome profiles across different samples and datasets. By scaling, transforming, or otherwise correcting the data, normalization methods help to harmonize microbiome datasets, making them amenable to accurate predictions of various phenotypic outcomes. However, the impact of normalization methods on predictions mainly focused on DNA microarray data and RNA-Seq data [9, 10]. Given the central role of normalization in microbiome data analysis and the lack of current methods comparison for microbiome data, there is a need to systematically evaluate their performance.

In our previous study [11], we evaluated existing normalization methods and assessed their efficacy in predicting binary phenotypes using microbiome data. Building on this, our current study examines the effects of heterogeneity on predicting quantitative phenotypes and aims to assess the performance of various normalization methods in predicting quantitative phenotypes across studies. To conduct this investigation, we utilized a diverse and extensive dataset comprising 31 shotgun sequencing datasets obtained from healthy stool samples. Each dataset was paired with a separate dataset for training and testing purposes separately, allowing for a thorough evaluation of prediction performance. We used the Root Mean Square Error (RMSE) as the primary performance metric, given its significance in quantifying prediction accuracy. Additionally, we supplemented our analysis with simulation studies that address three types of heterogeneity: background distributions of taxa in populations, batch effects across studies from the same population, and phenotype-associated models in different studies. These simulations enabled us to evaluate the performance of normalization methods in controlled settings, yielding valuable insights into how they perform under different scenarios.

This study aims to inform researchers on the necessary knowledge to make informed decisions when analyzing metagenomic data. Ultimately, this research aims to improve the reliability and accuracy of predictions obtained from metagenomic datasets, advancing our understanding of the complex relationships between microbial communities and host phenotypes.

## Materials and methods

### curatedMetagenomicData 3.8.0

The curatedMetagenomicData 3.8.0 package presents a curated meta-dataset of the human microbiome, derived from a collection of 93 cohorts involving shotgun sequencing of six distinct body sites. The raw sequencing data underwent a rigorous and standardized processing pipeline. Each sample in this dataset includes six primary data categories: gene family, marker abundance, marker presence, pathway abundance, pathway coverage, and relative taxonomic abundance values. Taxonomic abundance values were determined using MetaPhlAn3 [12], while the assessment of metabolic functional potential was performed through HUMAnN3 [13]. The package provides curated clinical and phenotypic metadata. For more comprehensive insights, please refer to the official documentation of the curatedMetagenomicData package [14].

In order to compare the predictive performance of different methods for normalizing microbiome profiles in predicting BMI values, our analysis focuses specifically on healthy subjects obtained from the curatedMetagenomicData dataset. We selected subjects from all cohorts based on the following inclusion criteria: (1) stool samples; (2) healthy status; (3) no missing BMI values; (4) read counts exceeding 1250. Additionally, if multiple samples were available for a subject, we randomly selected one sample. We only included datasets with a sample size greater than 30. In total, our analysis involved 5963 samples from 31 datasets. Table 1 presents the characteristics of the curatedMetagenomicData datasets used in our analysis.

**Table 1.** Characteristics of curatedMetagenomicData datasets involved in our analysis, with different DNA extraction kits (DNA-Exk) and sequencing platforms (Seq-Plat).

| Dataset | Country | Sample size | DNA-Exk | Seq-Plat | Reference |
| --- | --- | --- | --- | --- | --- |
| AsnicarF_2021 | USA/GBR | 1097 | PowerSoilPro | IlluminaNovaSeq | [15] |
| CosteaPI_2017 | KAZ | 84 | Gnome | IlluminaHiSeq | [16] |
| DeFilippisF_2019 | ITA | 97 | PowerSoil | IlluminaNextSeq | [17] |
| DhakanDB_2019 | IND | 107 | Qiagen | IlluminaNextSeq | [18] |
| HansenLBS_2018 | DNK | 58 | NA | IlluminaHiSeq | [19] |
| HMP_2012 | USA | 95 | Qiagen | IlluminaHiSeq | [20] |
| JieZ_2017 | CHN | 164 | Qiagen | IlluminaHiSeq | [21] |
| KarlssonFH_2013 | SWE/DEU/FRA/ISL | 43 | NA | IlluminaHiSeq | [22] |
| KaurK_2020 | IND | 31 | ZR_Fecal_DNA_MiniPrep | IlluminaHiSeq | [23] |
| KeohaneDM_2020 | IRL | 116 | NA | IlluminaHiSeq | [24] |
| LeChatelierE_2013 | DNK | 115 | NA | IlluminaHiSeq | [25] |
| LifeLinesDeep_2016 | NLD | 1135 | Qiagen | IlluminaHiSeq | [26] |
| LokmerA_2019 | CMR | 56 | Illuminakit | IlluminaHiSeq | [27] |
| NagySzakalD_2017 | USA | 50 | KAMA_Hyper_Prep | IlluminaHiSeq | [28] |
| NielsenHB_2014 | ESP | 59 | NA | IlluminaHiSeq | [29] |
| Obregon-TitoAJ_2015 | PER/USA | 51 | MoBio | IlluminaHiSeq | [30] |
| PasolliE_2019 | MDG | 112 | Qiagen | IlluminaHiSeq | [31] |
| QinJ_2012 | CHN | 174 | NA | IlluminaHiSeq | [32] |
| QinN_2014 | CHN | 114 | NA | IlluminaHiSeq | [33] |
| RubelMA_2020 | CMR | 86 | PSP_Spin_Stool | IlluminaHiSeq | [34] |
| SchirmerM_2016 | NLD | 456 | Illuminakit | IlluminaHiSeq | [35] |
| ThomasAM_2018 | ITA | 39 | Qiagen/Gnome | IlluminaHiSeq | [3] |
| VogtmannE_2016 | USA | 52 | Gnome | IlluminaHiSeq | [36] |
| WirbelJ_2018 | DEU | 65 | Gnome | IlluminaHiSeq | [6] |
| XieH_2016 | GBR | 169 | Qiagen | IlluminaHiSeq | [37] |
| YachidaS_2019 | JPN | 245 | NA | IlluminaHiSeq | [38] |
| YeZ_2018 | CHN | 45 | Qiagen | IlluminaHiSeq | [39] |
| YuJ_2015 | CHN | 38 | Qiagen | IlluminaHiSeq | [40] |
| ZeeviD_2015 | ISR | 870 | NA | IlluminaHiSeq | [41] |
| ZellerG_2014 | FRA | 59 | Gnome | IlluminaHiSeq | [42] |
| ZhuF_2020 | CHN | 81 | NA | IlluminaHiSeq | [43] |

### Statistical Methods

We performed microbial relative abundance calculations for each sample and computed the Shannon indices using the *diversity()* function from the R package *vegan* [44]. The differences in Shannon indices between each dataset and the overall Shannon indices were determined using the Wilcoxon rank sum test. The dissimilarities between sample pairs were quantified using the Bray-Curtis distance [45], implemented by the *vegdist()* function from the R package *vegan* [44]. Principal coordinate analysis (PCoA) was employed to effectively visualize the sample clustering, using the *pcoa()* function from the R package *ape* [46]. To assess the variance attributable to population factors, we conducted permutational multivariate analysis of variance (PERMANOVA) [47] using the *adonis()* function in the R package *vegan* [44]. The total variance explained by each variable was evaluated independently of other variables, to avoid issues with variable ordering, and thus should be regarded as the total variance explainable by that variable [48].

### Normalization Methods

Microbiome data analysis commonly employs a range of normalization methods. In predicting the quantitative traits of unknown samples, it is crucial to transform or normalize the data to ensure that both the training and testing datasets adhere to the same underlying distribution. This investigation encompassed a comprehensive comparative analysis, examining seven scaling methods, one approach based on compositional data analysis, eight transformation methods, and six batch correction methods. To the best of our knowledge, this study represents the most thorough comparison conducted to date, focused on prediction.

The taxa count table of a metagenomic dataset can be structured as shown in Table 2. Suppose we have a dataset with *n* samples and *m* features. Denote the count for taxon *i* in sample *j* as *c*_*ij*_. With this notation, the procedures and equations for normalization methods can be outlined as follows.

**Table 2.** Format of a count table of metagenomic dataset, where *c*_*ij*_ is the number of reads belonging to taxon *i* and sample *j*.

| | sample 1 | ... | sample $j$ | ... | sample $n$ |
| --- | --- | --- | --- | --- | --- |
| taxon 1 | $c_{11}$ | ... | $c_{1j}$ | ... | $c_{1n}$ |
| ⋮ | ⋮ | ... | ⋮ | ... | ⋮ |
| taxon $i$ | $c_{i1}$ | ... | $c_{ij}$ | ... | $c_{in}$ |
| ⋮ | ⋮ | ... | ⋮ | ... | ⋮ |
| taxon $m$ | $c_{m1}$ | ... | $c_{mj}$ | ... | $c_{mn}$ |

### Scaling Methods

Scaling is a commonly utilized method for normalizing microbiome data, with the primary objective of reducing biases introduced by sequencing technology. This is achieved by dividing the counts in the taxa count table by a scaling or normalization factor. Mathematically, this can be represented by the following equation:

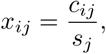

where *x*_*ij*_ is the normalized abundance for taxon *i* in sample *j*, and *s*_*j*_ is the scaling/normalization factor for sample *j*. We investigated seven popular scaling methods (Table 3) in our analysis, including TSS, UQ, MED, CSS in *metagenomeSeq*, TMM in *edgeR*, RLE in *DESeq2*, and GMPR in *GUniFrac*.

**Table 3.**
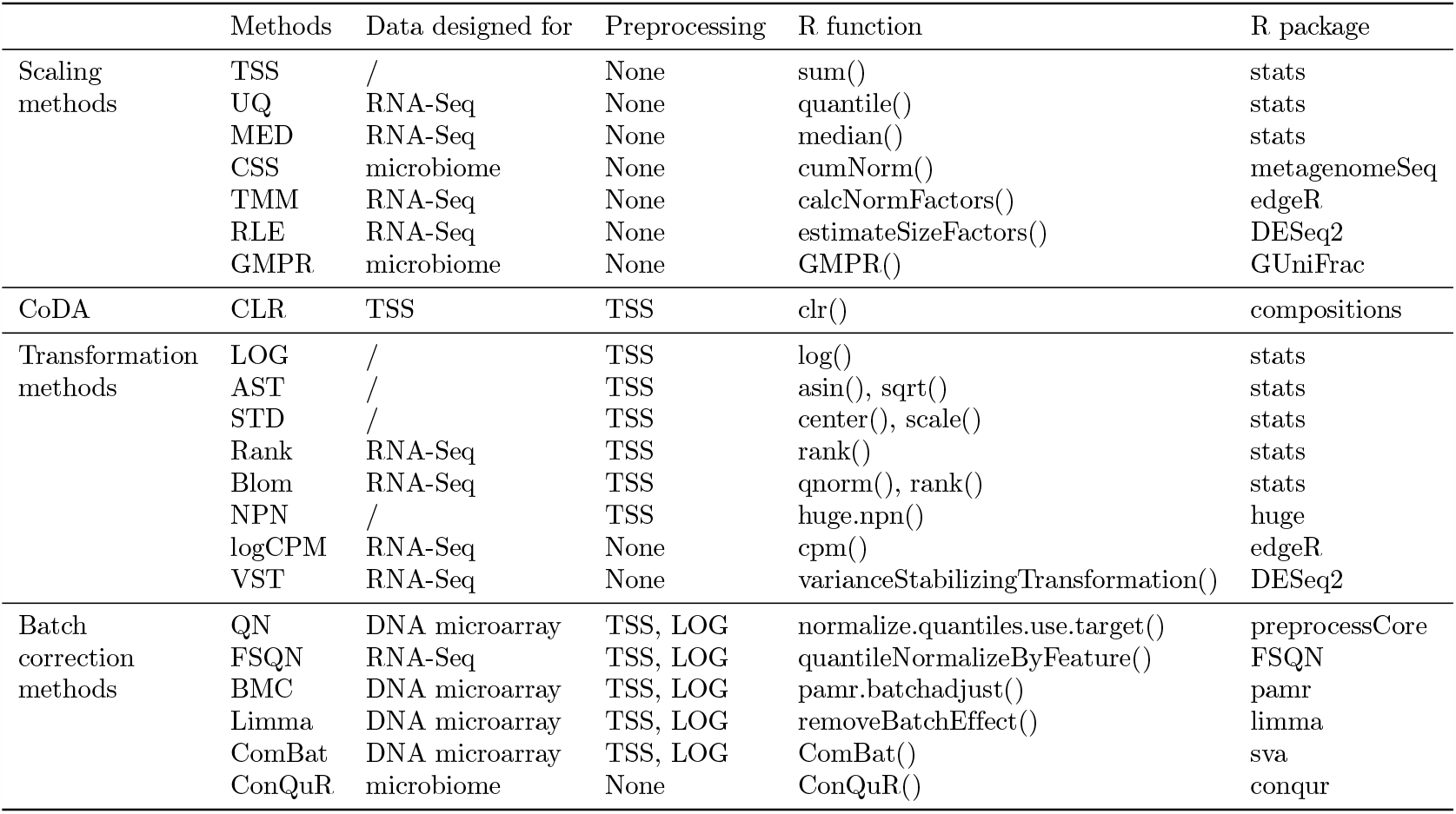
Summary of normalization methods, including seven scaling methods, one compositional data analysis (CoDA) method, eight transformation methods, and six batch correction methods.

### Total Sum Scaling (TSS) [49]

The TSS method divides counts by the total number of reads in a sample. This can be expressed by the equation:

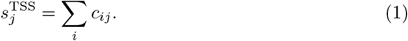

### Upper Quartile (UQ) [49, 50]

The UQ method scales each sample by the upper quartile of counts that are different from zero in that sample. This can be expressed by the equation:

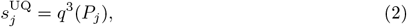

where *q*^3^(·) represents the function of estimating upper quartile, and

*P*_***j***_ ={ ***c***_***ij***_|***c***_***ij***_ ***>*** 0, ***i* =** 1, · · ·, *n*} represents a set of counts different from 0 in sample *j*.

### Median (MED) [49]

The MED method calculates the scaling factor based on the median of counts that are different from zero. It can be expressed as follows:

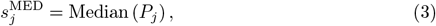

where Median(*·*) is the function that estimates the median, and

*P*_*j*_ ={*c*_*ij*_ |*c*_*ij*_ *>* 0, *i* = 1, · · ·, *n*} represents a set of counts different from zero in sample *j*.

### Cumulative Sum Scaling (CSS) [51]

CSS modified TSS for microbiome data on a sample-specific basis. It determines the scaling factor as the cumulative sum of counts, up to a percentile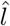 determined by the data. The scaling factor can be calculated using the following equation:

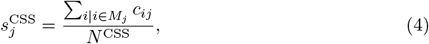

Where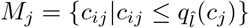 represents the taxa included in the cumulative summation for sample *j*, and *N* ^CSS^ is an appropriately chosen normalization constant. This scaling method is implemented using the *cumNorm()* function in the R package *metagenomeSeq* [51].

### Trimmed Mean of M-values (TMM) [52]

TMM is a widely used normalization method for RNA-Seq data, assuming that the majority of genes are not differentially expressed. It operates by selecting a reference sample and treating the other samples as test samples. If not specified, the reference sample is chosen as the one with the count-per-million upper quantile closest to the mean upper quantile. The scaling factor between a test sample and the reference sample is estimated based on the ratio of two observed relative abundances, known as M values and A values. Specifically, the *M* value, denoted as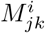, is defined as the log2 ratio of the taxon *i* in sample *j* to its abundance in sample *k*. On the other hand, the *A* value, denoted as 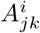, is the log2 of the geometric mean of the observed relative abundances. By default, the TMM method trims the *M* values by 30% and the *A* values by 5%. The scale factor 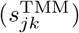of sample *j* to sample *k* can then be calculated as the weighted sum of the *M* values:

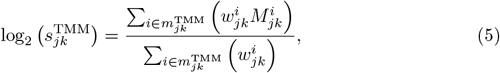

Where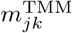 denotes the remaining taxa after the trimming step, and weight 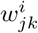is given by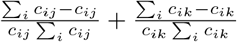 . The TMM scaling method can be implemented using *calcNormFactors()* function in the *edgeR* [53] Bioconductor package.

### Relative log expression (RLE) [54]

RLE is another widely used method for RNA-Seq data and relies on the same assumption that there is a large invariant part in the count data. It first calculates the geometric mean of the counts to a gene from all the samples and then computes the ratio of a raw count over the geometric mean to the same gene. The scale factor for a particular sample is obtained as the median of these ratios:

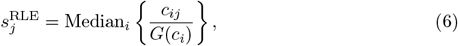

Where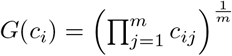 is the geometric mean of gene *i*. To account for zeros in microbiome data, a modified geometric mean is computed by taking the n-th root of the product of the non-zero counts. This calculation is performed by setting the *type=“poscounts”* option in the *estimateSizeFactors()* function of the *DESeq2* [55] Bioconductor package.

### Geometric mean of pairwise ratios (GMPR) [56]

GMPR extends the idea of RLE normalization by reversing the order of computing geometric and median. This modification aims to address the issue of zero inflation that occurs in microbiome data. The scale factor for a given sample *j* using reference sample *k* is determined by the following equation:

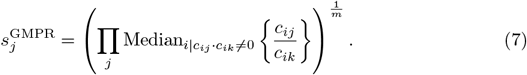

This scaling method is implemented using *GMPR()* function in the *GUniFrac* [57] package.

### Compositional Data Analysis (CoDA) Methods

Gloor et al. [58] emphasized that microbiome datasets generated through high-throughput sequencing are inherently compositional due to the arbitrary total imposed by the sequencing instrument. Consequently, various approaches have been proposed to mitigate the impact of sampling fractions by converting the abundances into log ratios within each sample. Commonly employed methods in compositional data analysis include the Additive Log-Ratio transformation (ALR) [59], the Centered Log-Ratio transformation (CLR) [59], and the Isometric Log-Ratio transformation (ILR) [59]. ALR and ILR transform an *n*-dimensional gene vector into an (*n−*1)-dimensional dataset in Euclidean space, requiring the selection of a reference gene. Due to computational challenges arising from the large number of genes, our analysis solely focused on CLR.

### Centered Log-Ratio (CLR) [59]

CLR transformation is a compositional data transformation that calculates the log-ratio of counts and their geometric means within each sample, using relative abundances. Mathematically, this transformation can be represented as:

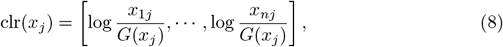

where *x*_*ij*_ is the relative abundance of gene *i, i* = 1, · · ·, *n* in sample *j, j* = 1, · · ·, *m*,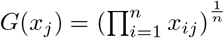 is the geometric mean of sample *j* with a pseudo count 0.65 times minimum non-zero abundance added to 0 values [60]. The CLR transformation can be implemented using *clr()* function in R package *compositions* [61].

### Transformation Methods

Microbiome data exhibit several problematic properties, including skewed distributions, unequal variances for individual taxa, and extreme values. To address these issues when fitting the prediction model, we propose applying transformations to the microbiome data. These transformations can be used to address one, two, or all of these problems. Specifically, let *c*_*ij*_ and *x*_*ij*_ represent the count and relative abundance of gene *i, i* = 1, · · ·, *n* in sample *j, j* = 1, · · ·, *m*. In this study, we explore various transformation methods, as outlined in Table 3. These methods include LOG, AST, STD, Rank, Blom, NPN in *huge*, logCPM in *edgeR*, and VST in *DESeq2*.

### LOG

Log transformation is frequently employed to address taxa with skewed distributions, thereby achieving transformed abundances that are closer to a normal distribution [9]. To prevent infinite values, a pseudo count of 0.65 times the minimum non-zero abundance is added to the zero values before applying the log transformation [60].

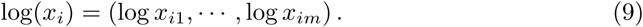

### Arcsine square-root (AST)

AST transformation is used to reduce the presence of extreme values in the data and achieve a more approximately normal distribution. Mathematically, it is defined as:

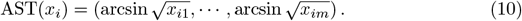

### Standardization (STD) [9]

STD is the default implementation in many regression analyses to reduce the variations of features (taxa in our analysis). Mathematically, it is defined as:

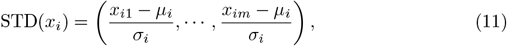

where *µ*_*i*_ and *σ*_*i*_ is the mean and standard deviation of gene *i* separately.

### Rank [9]

Rank transformation is a widely used and straightforward method in non-parametric statistics. The features that undergo rank transformation are distributed uniformly between zero and the sample size *m*. A small noise term *ϵ*_*ij*_ *∼ N* (0, 10^*−*10^) is added before data transformation to handle the ties of zero counts.

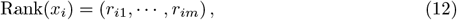

Rank(*x*_*i*_) = (*r*_*i*1_, · · ·, *r*_*im*_), (12) where *r*_*ij*_, *j* = 1, *· · ·, m* is the corresponding rank for relative abundance *x*_*ij*_, *j* = 1, · · ·, *m* in gene *i*.

### Blom [9, 62]

Blom transformation is based on rank transformation. The uniformly distributed ranks are further transformed into a standard normal distribution using the following equation:

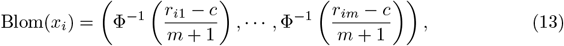

Where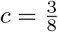 is a constant, Φ^*−*1^(·) denotes the quantile function of normal distribution, and *r*_*ij*_, *j* = 1, · · ·, *m* is the corresponding rank for relative abundance *x*_*ij*_, *j* = 1, · · ·, *m* in gene *i*.

### Non-paranormal (NPN) [63]

The NPN transformation is intended to be incorporated into an enhanced graphical lasso approach, which initially converts variables into univariate smooth functions to estimate a Gaussian copula. However, this transformation can also be used independently for analysis purposes. By denoting the Gaussian cumulative distribution function as Φ, we can estimate the transformed data by

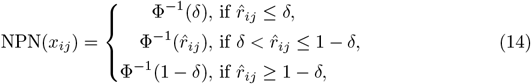

Where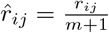 and 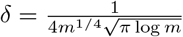. The implementation of this transformation can be performed using the *huge*.*npn()* function in R package *huge* [64].

### Log counts per million (logCPM)

The logCPM transformation represents the logarithm of counts per million, which serves as a descriptive measure for the gene expression level in RNA-Seq data. We employed this transformation on the microbiome data by adding a pseudo count equal to 0.65 times the minimum non-zero abundance to the zero values prior to the logarithmic transformation. The logCPM transformation for gene *i* can be mathematically expressed as

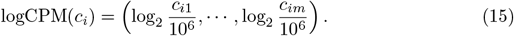

This transformation method is implemented using *cpm()* function in the *edgeR* [53] Bioconductor package.

### Variance Stabilizing Transformation (VST) [54]

VST models the relationship between mean *µ*_*i*_ and variance 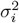for each gene *i*:

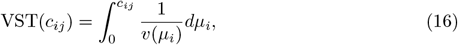

where 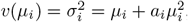, with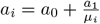 being a dispersion parameter and *a*_0_ and *a*_1_ are estimated in a generalized linear model. A pseudo count 1 was added to zero values. This transformation is implemented using *varianceStabilizingTransformation()* function in the *DESeq2* [55] Bioconductor package.

### Batch Correction Methods

Batch effects frequently occur in genomic technologies and can result from various specimen processing steps. However, normalization methods alone may not adequately address these batch effects. Consequently, multiple approaches have been proposed to effectively remove batch effects. In this study, we examined six commonly used methods: QN implemented in the *preprocessCore* package, FSQN implemented in *FSQN*, BMC implemented in *pamr*, Limma implemented in *limma*, ComBat implemented in *sva*, and ConQuR implemented in *conqur* (Table 3).

### Quantile normalization (QN) [65]

QN was initially developed for DNA microarrays, but has since been adapted for various data types, including microbiome data. This method replaces each value in a target distribution with the corresponding value from a reference distribution based on their rank order. In cases where the reference distribution encompasses multiple samples, the reference distribution should be first quantile normalized across all samples [66]. In our analysis, used the training data as the reference distribution. We applied QN to log-transformed relative abundances, substituting zeros with a pseudo count that was calculated as 0.65 times the minimum non-zero abundance across the entire abundance table. The reference distribution is obtained using function *normalize*.*quantiles*.*determine*.*target()* in R package *preprocessCore* [67]. And the batch effects are removed using function *normalize*.*quantiles*.*use*.*target()* in R package *preprocessCore* [67].

### Feature specific quantile normalization (FSQN) [10]

FSQN is a variation of quantile normalization (QN) that normalizes genes instead of samples. In FSQN, the reference distribution consists of the genes in the training set, while the target distribution consists of the genes in the testing set. It is applied to log-transformed relative abundance data, with zeros replaced with pseudo count 0.65 times the minimum non-zero abundance across the entire abundance table, using function *quantileNormalizeByFeature()* in R package *FSQN* [10].

### Batch mean centering (BMC) [68]

BMC is a method that centers the data on a batch-by-batch basis. The mean abundance per gene for a given dataset is subtracted from the individual gene abundance. It is applied to log-transformed relative abundance data, with zeros replaced with pseudo count 0.65 times the minimum non-zero abundance across the entire abundance table. To perform BMC, the function *pamr*.*batchadjust()* from the R package *pamr* [69] is utilized.

### Linear models for microarray data (Limma) [70]

Limma is a method that utilizes linear modeling to remove batch effects. We first calculate the relative abundances and apply a log2 transformation to them. A pseudo count 0.65 times the minimum non-zero abundance across the entire abundance table was added to zeros to avoid infinite values for log transformation. The *removeBatchEffect()* function in R package *limma* [70] is then used to correct for batch effects.

### ComBat [71]

ComBat employs an empirical Bayes framework to estimate and remove batch effects while preserving the biological variation of interest. Before batch correction, the relative abundance of microbiome data was log-transformed. Zeros were replaced with a pseudo count calculated as 0.65 times the minimum non-zero abundance across the entire abundance table. This correction method is implemented in R using the *ComBat()* function from the *sva* package.

### Conditional quantile regression (ConQuR) [72]

ConQuR is employed to remove batch effects from a count table. This batch correction method is implemented in R using the ConQuR function from the ConQuR package [72].

## The Random Forest classifiers

The random forest algorithm is a supervised learning approach that is capable of handling both regression and classification problems [73]. It has been shown to outperform other machine learning tools when applied to microbiome data [74]. In this study, we utilized random forest regression to predict a quantitative phenotype. The implementation was carried out using the *train()* function from the R package *caret* [75]. We constructed a random forest with 1,000 decision trees, and the number of variables at each decision tree was optimized through grid search using 10-fold cross-validation.

To ensure the comparability and integrity of our training and testing datasets, we applied normalization procedures aimed at reducing heterogeneity within and across datasets. For scaling methods that involve selecting references, such as TMM and RLE, and transformation methods that ensure prediction covariates (taxa) are drawn from the same distribution, such as STD, Rank, Blom, NPN, and VST, we performed normalization on the training data. We then applied an additional normalization step to the test data based on the training data. This approach guarantees that the normalization of the training data remains independent of the testing data [76].

We evaluated the performance of our predictions using the Root Mean Squared Error (RMSE), which quantifies the square root of the average squared differences between predicted and actual values. The RMSE is defined as follows:

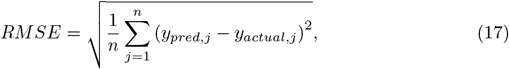

where *n* is the number of observations, *y*_*pred,j*_ is the predicted value for sample *j, y*_*actual,j*_ is the actual value for sample *j*.

## Simulation Study

In line with our previous investigation on case-control studies [11], we devised three unique scenarios to account for the heterogeneity within the training and testing data. For each combination of the parameters, we iterated the procedure 100 times. Subsequently, the datasets underwent normalization using various methods. Employing the random forest algorithm, we constructed prediction models based on one simulated population and evaluated their performance on the other population in each of the three scenarios. To assess the accuracy of the predictions, we computed the Root Mean Squared Error (RMSE) values for the 100 simulation runs conducted across the different scenarios.

### Scenario 1: Different background distributions of taxa in populations

In the first scenario, we considered that the variations between populations were attributable to differences in the underlying distributions of taxa, such as ethnicity or diet. McMurdie and Holmes [77] proposed a method to simulate samples from distinct populations (Simulation A) and samples with case-control designs (Simulation B) independently within this particular scenario. In our simulations, we combined these strategies and implemented specific modifications.

Our methodology commenced by establishing the baseline levels of taxon abundance for the training and testing populations. To replicate this scenario, we procured two publicly available and geographically diverse datasets using the control samples from GuptaA 2019 and FengQ 2015 as the template for our simulations, which is the same with our previous analysis [11]. Specifically, we included 30 control samples and 183 species from the GuptaA 2019 dataset [18, 78] for training purposes, and 61 healthy samples and 468 species from the FengQ 2015 dataset [79] for testing purposes. For each dataset, we were provided with a count table consisting of rows representing taxa and columns representing samples. By summing the rows, we obtained the initial vectors representing the underlying taxon abundance in different populations, denoted as *p*_*k*_, where *k* = 1, 2.

To explore the influence of dissimilarities between two populations on cross-study prediction, we constructed pseudo-population vectors *v*_*k*_, where *k* = 1, 2:

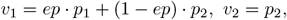

where *ep* denotes the population effect that quantifies the differences between two populations. Importantly, it should be emphasized that *v*_1_*′ −v*_2_*′* = *ep*(*v*_1_*− v*_2_), which underscores how the differences between the two simulated populations escalate with increasing values of *ep*. By incrementally varying *ep* from 0 to 1 in intervals of 0.2, we analyzed the overall trends of different normalization methods.

To generate pseudo read counts for 100 samples within each population, we posited that the taxonomic probabilities *x*_*kj*_ of sample *j* belonging to population *k* followed a Dirichlet distribution *Dir*(*α*_*k*_), with *α*_*k*_ = *c · v*_*k*_ for *k* = 1, 2. To ensure minimal variation, we assigned a large value to *c*, resulting in a variance of *x*_*kj*_ that approximates 0 and aligns with *v*_*k*_. To introduce some level of variability, we selected *c* = 1 *×* 10^6^ (preventing the generation of zero probabilities). The read counts for each sample were simulated using a multinomial distribution *MN* (library size, *x*_*kj*_), *k* = 1, 2, where the library size was set to 1, 000, 000 and the probabilities were derived from the Dirichlet distribution.

Among the 154 taxa shared by the two populations, we randomly chose 10 taxa and proposed that these taxa were linked to a specific quantitative phenotype of interest. It was assumed that the first 5 taxa exhibited enrichment while the remaining 5 were diminished. A vector of pseudo coefficients was generated from a uniform distribution with lower and upper bounds of 3 and 5 for positive associations, and*−* 5 and *−* 3 for negative associations. The chosen taxa and their corresponding pseudo coefficients remained consistent throughout the simulations. The quantitative phenotypes were simulated based on the relationship between the phenotype and the corresponding microbial abundances as follows:

- Linear: *y* = *c*_1_ * *β*^*T*^ *x* + ϵ
- Quadratic: *y* = *c*_2_ * *β*^*T*^ *x*^2^ ϵ
- Inverse: 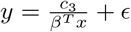
- Logistic: 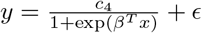

where *x* is the vector of the selected phenotype associated with microbial relative abundance, *β* indicates the pseudo coefficients, *c*_1_, *c*_2_, *c*_3_, *c*_4_ represent constants used to control the range of absolute values of the simulated phenotypes *y* (ranging in dozens or hundreds), and *ϵ ∼ N* (0, 1) represents random noise.

### Scenario 2: Different batch effects of studies with the same background distribution of taxa in populations

In this scenario, we employed the controls in FengQ 2015 dataset [79] as the template for our simulations, ensuring that the background distribution remained consistent between the training and testing datasets. By doing so, we effectively eliminated the population effects observed in Scenario 1. The generation of read counts and phenotypes followed the same procedure as in Scenario 1, utilizing multinomial distributions with a sample size of one million reads. Specifically, we specified 10 taxa associated with the phenotype, and considered linear, quadratic, inverse, and logistic relationships between the phenotype and the corresponding microbial abundances.

To simulate batch effects, we followed a similar procedure as described in Zhang et al. [8]. We assumed that the mean (*γ*_*ik*_) and variance (*δ*_*ik*_) of taxon *i* were influenced by the batch *k*. Drawing from the batch effect generating model proposed by Johnson et al. [71], we assumed an additive effect on the mean and a multiplicative effect on the variance for each taxon. The values of *γ*_*ik*_ and *δ*_*ik*_ were randomly sampled from normal and inverse gamma distributions, respectively, as expressed by:

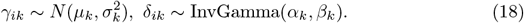

To specify the hyperparameters (*µ*_*k*_, *σ*_*k*_, *α*_*k*_, *β*_*k*_), we defined two values to indicate the severity of batch effects. Specifically, we considered three levels for the batch effect on the mean (*sev*_*mean*_ *∈ {*0, 500, 1000*}*) and three levels for the batch effect on the variance (*sev*_*var*_ *∈*{1, 2, 4}). For a given severity level, the variance of *γ*_*ik*_ and *δ*_*ik*_ was fixed at 0.01, while the batch effect parameters were either added or multiplied to the mean and variance of the original study’s expression. Importantly, the batch effects were solely applied to the training data, while the test dataset remained unaltered.

### Scenario 3: Different phenotype models of studies with the same background distribution of taxa in populations

In this scenario, we hypothesized that the model for phenotype-associated taxa may differ between populations. To mitigate the population effects mentioned in Scenario 1, we employed the FengQ 2015 dataset [79] as the template for simulations. In order to alleviate the batch effects described in Scenario 2, this simulation scenario did not incorporate any batch effects.

To select phenotype-associated taxa, we predetermined 10 taxa for the training data. From the initial 10 taxa, we selected a subset and added additional taxa to maintain a total of 10 signature taxa in the testing data. The level of resemblance between the training and testing data was determined by the number of taxa that overlapped, ranging from 2 to 10 with increments of 2. Subsequently, the two populations were simulated following the same procedure as in the previous two scenarios. The simulation parameters consisted of 100 samples per population, one million reads per sample, and four distinct relationships between quantitative phenotype and phenotype-associated taxa.

## Results

### The reproducibility of cross-study prediction is limited by various confounding variables

Using the inclusion criteria outlined in the curatedMetagenomicData 3.8.0 section, we incorporated a total of 5963 healthy stool samples into our analysis. These samples were obtained from 31 different datasets and exhibited various biological and technical differences, encompassing variations in geographic origin, DNA extraction techniques, and sequencing platforms (Table 1).

Initially, we examined the BMI values and Shannon indices within each dataset to identify the overall patterns in sample characteristics. S1 Fig demonstrates noticeable differences in BMI among samples from different datasets, with each dataset having its own distinct range. The average BMI values for each dataset varied significantly, ranging from 21.2 (DhakanDB 2019) to 30.7 (KeohaneDM 2020), while the overall average BMI for all samples was 24.9. To assess the significance of these variations, we performed Wilcoxon tests comparing the BMI of each dataset with the overall sample mean. Among the 31 datasets, 10 displayed significantly higher BMIs than the overall sample mean, while 11 exhibited significantly lower BMIs. The Shannon indices, as presented in S2 Fig, generally mirrored the trends observed in BMI values, although differences persisted. Notably, KeohaneDM 2020, despite having the highest average BMI, demonstrated significantly lower Shannon indices compared to the overall dataset. Conversely, LeChatelierE 2013 displayed a significantly higher average BMI than the overall average but exhibited no significant differences in Shannon indices.

The similarities among different datasets were evaluated by generating a PCoA plot using the Bray-Curtis distance metric, as depicted in Fig 1A. Owing to the large sample sizes, the mean point of each dataset was used to represent the positions of the samples from that dataset on the PCoA plot. Additionally, the size of the points indicated the sample size of each dataset. This plot revealed distinct separations between the datasets, indicating variations in microbial composition. To further assess the contribution of biological or technical factors to microbiome variation, PERMANOVA was performed on the Bray-Curtis distance. Fig 1B illustrates that all seven factors considered in the analysis accounted for a significant proportion of the variations. The three most influential factors affecting the community structures of the microbiome data were datasets, country, and DNA extraction kit, followed by sequencing platform, age, gender, and BMI.

**Fig 1.**
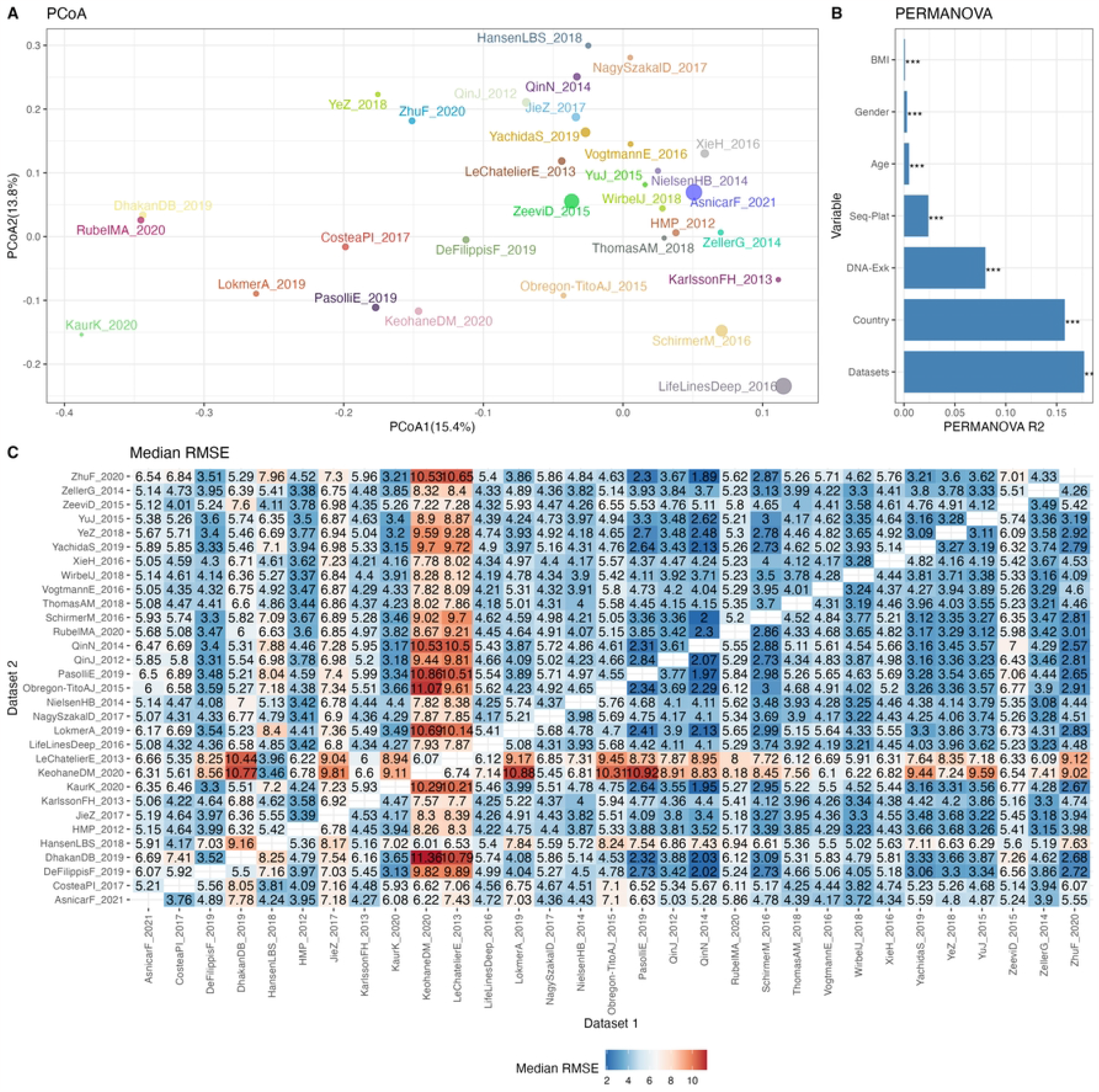
The reproducibility of cross-datasets prediction is limited by various confounding variables. A: PCoA plot based on Bray-Curtis distance, with colors for different datasets. B: Bar plot of variance explained by different variables (R2) from PERMANOVA based on Bray-Curtis distance. P values calculated by 1,000 permutations were annotated on the top of the bar, with ^***^ for *p <* 0.001. C: Median RMSE of cross-datasets prediction based on TSS normalized abundances using random forests model over 10 repetitions.

Subsequently, the impact of heterogeneities on the reproducibility of BMI prediction was examined. The classifier was trained on each individual training set (dataset 1), and the model was applied to each testing set (dataset 2). The cross-prediction matrix of RMSE values, obtained using random forest models on TSS normalized abundances, is illustrated in Fig 1C. The median RMSE values exhibited variation across datasets and were influenced by multiple factors. Importantly, a significant disparity in RMSE values was observed when different datasets were used as the training dataset to predict KeohaneDM 2020. Specifically, 18 out of 30 datasets exhibited RMSE values exceeding 7, indicating an inaccurate prediction of the BMI in KeohaneDM 2020. This finding aligned with the substantial differences in microbiome composition compared to the other datasets. LeChatelierE 2013 was another dataset demonstrating relatively poor prediction repeatability, despite not differing significantly from other datasets in terms of its Shannon indices. In contrast, the dataset LifeLinesDeep 2016 displays significant differences in the PCoA plot compared to other datasets, yet it performs well in predicting and can also be effectively predicted by other datasets. This phenomenon can possibly be attributed to its relatively larger sample size. In conclusion, these results indicate that the reproducibility of response predictions were influenced by various factors.

The presence of confounding factors, such as country and DNA extraction kit, led to notable variations in the background distributions of taxa. We conducted an evaluation to ascertain whether models trained on one dataset could accurately predict a quantitative phenotype for samples in another dataset. Additionally, we examined whether the implementation of normalization methods could enhance prediction performance. To achieve this, we employed seven scaling methods, one compositional data analysis method, eight transformation methods, and six batch correction methods on the simulated training and testing data.

### Batch correction methods are necessary for quantitative phenotype prediction

In our simulation studies, we evaluate the influence of heterogeneity on prediction performance across three distinct scenarios and using four different types of quantitative phenotypes. Additionally, we analyze and compare the prediction performance of various normalization methods.

In Scenario 1, we evaluate the impact of various normalization methods on the prediction of quantitative phenotypes across diverse background distributions of taxa. The experiments are repeated 100 times, and the median RMSE is calculated based on abundance profiles normalized by different methods. The results are presented in Fig 2. We find that prediction accuracy increases as population effects increase, as evident from the corresponding increase in RMSE values. However, different normalization methods exhibit minimal variation in predicting quantitative phenotypes. For instance, considering the linear relationship of phenotypes (Fig 2A), at a population effect of 0.2, the maximum and minimum RMSE values for different methods differ by 0.08. This difference slightly increases with an increase in population effect, peaking at a population effect of 1, where it reaches 0.23 - still a very small difference. Similar trends are observed for quadratic (Fig 2B), inverse (Fig 2C), and logistic (Fig 2D) relationships of phenotypes. These findings suggest that, among the 22 normalization methods we compared, none significantly outperforms the others in predicting quantitative phenotypes when the population effect is fixed.

**Fig 2.**
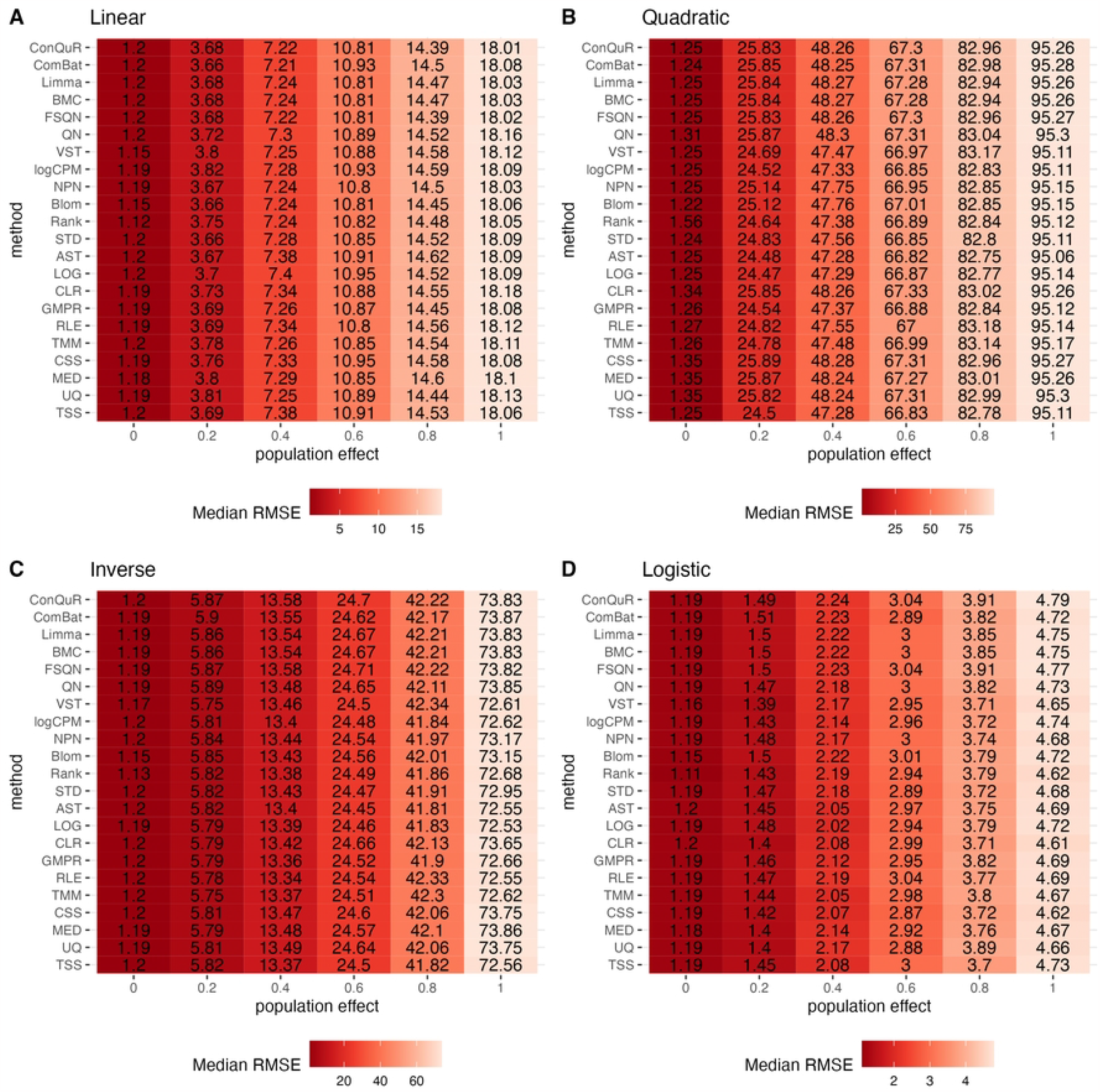
Heatmaps depicting median RMSE values obtained from abundance profiles normalized by different methods for predicting simulated quantitative phenotype in Scenario 1. The panels correspond to relationships between phenotype and phenotype-associated taxa, including A: linear, B: quadratic, C: inverse, and D: logistic. The columns represent different values of population effects, while the rows represent different normalization methods

In Scenario 2, we investigate the impact of batch effects on model prediction performance when utilizing abundance profiles normalized by different methods. Fig 3 demonstrates the median RMSE obtained from random forest models using abundance profiles normalized by different methods across 100 runs. As expected, we observe an increasing trend in RMSE values for all methods as batch effects increase. Interestingly, we find that batch effects of the mean taxa abundance appeared to have a greater impact than the variance of taxa abundance, especially in terms of model prediction performance. Furthermore, when compared to scaling methods and transformation methods, batch correction methods exhibit lower RMSE values, particularly when large batch mean differences are present. Among the six batch correction methods, in the case of a logistic relationship of phenotypes (Fig 3D), when the batch variance is set to 1 and the batch mean to 500 or 1000, BMC and Limma demonstrate similar performance, with the lowest predictive accuracy among all normalization methods. However, their respective RMSE values differ by no more than 0.05 from the minimum RMSE value. This minimal difference can be considered negligible in practical predictions. Additionally, the performance of ComBat is noteworthy. It exhibits lower predictive accuracy than other batch correction methods in linear (Fig 3A), quadratic (Fig 3B), and inverse (Fig 3C) relationships of phenotypes. However, in the case of a logistic relationship of phenotypes, it outperforms all other methods. This inconsistency highlights the need for caution when using ComBat for batch correction.

**Fig 3.**
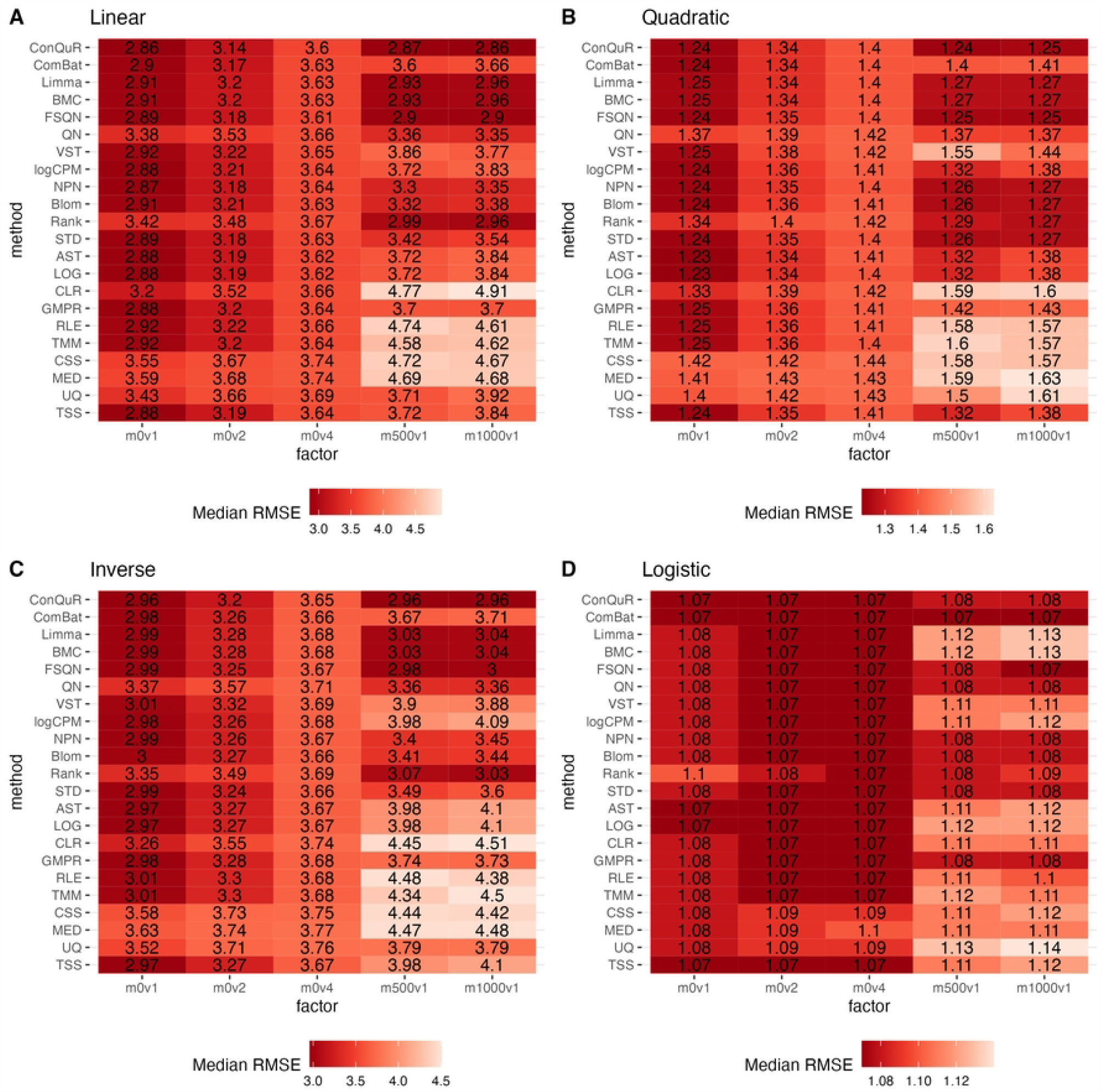
Heatmaps depicting median RMSE values obtained from abundance profiles normalized by different methods for predicting simulated quantitative phenotype in Scenario 2. The panels correspond to relationships between phenotype and phenotype-associated taxa, including A: linear, B: quadratic, C: inverse, and D: logistic. The columns represent different combinations of batch mean and batch variation, with “m” for batch mean adjusting the mean and “v” for batch variance adjusting the variance. The rows represent different normalization methods.

Fig 4 illustrates the findings from simulation scenario 3, which investigates the influence of different phenotype-associated feature models. Ideally, as the number of overlapping phenotype-associated taxa increases, the RMSE values should decrease. However, the choice of these taxa can significantly impact the prediction of quantitative phenotypes due to the shared background distributions of taxa. If randomly selected phenotype-associated taxa predominantly have zero values, it leads to similarity in the phenotype model during training and testing. This phenomenon is particularly noticeable when the overlapping number is 2 in both linear (Fig 4A) and quadratic (Fig 4B) relationships of phenotypes. In these cases, the median RMSE value at overlap=2 is lower than the median RMSE value at overlap=4. Across the four different types of quantitative phenotypes, the RMSE reaches its minimum at overlap=10, suggesting that at this point, phenotypes can be accurately predicted. However, similar to scenario 1, the performance of different normalization methods remains relatively consistent, with no single method significantly outperforming the others in predicting quantitative phenotypes.

**Fig 4.**
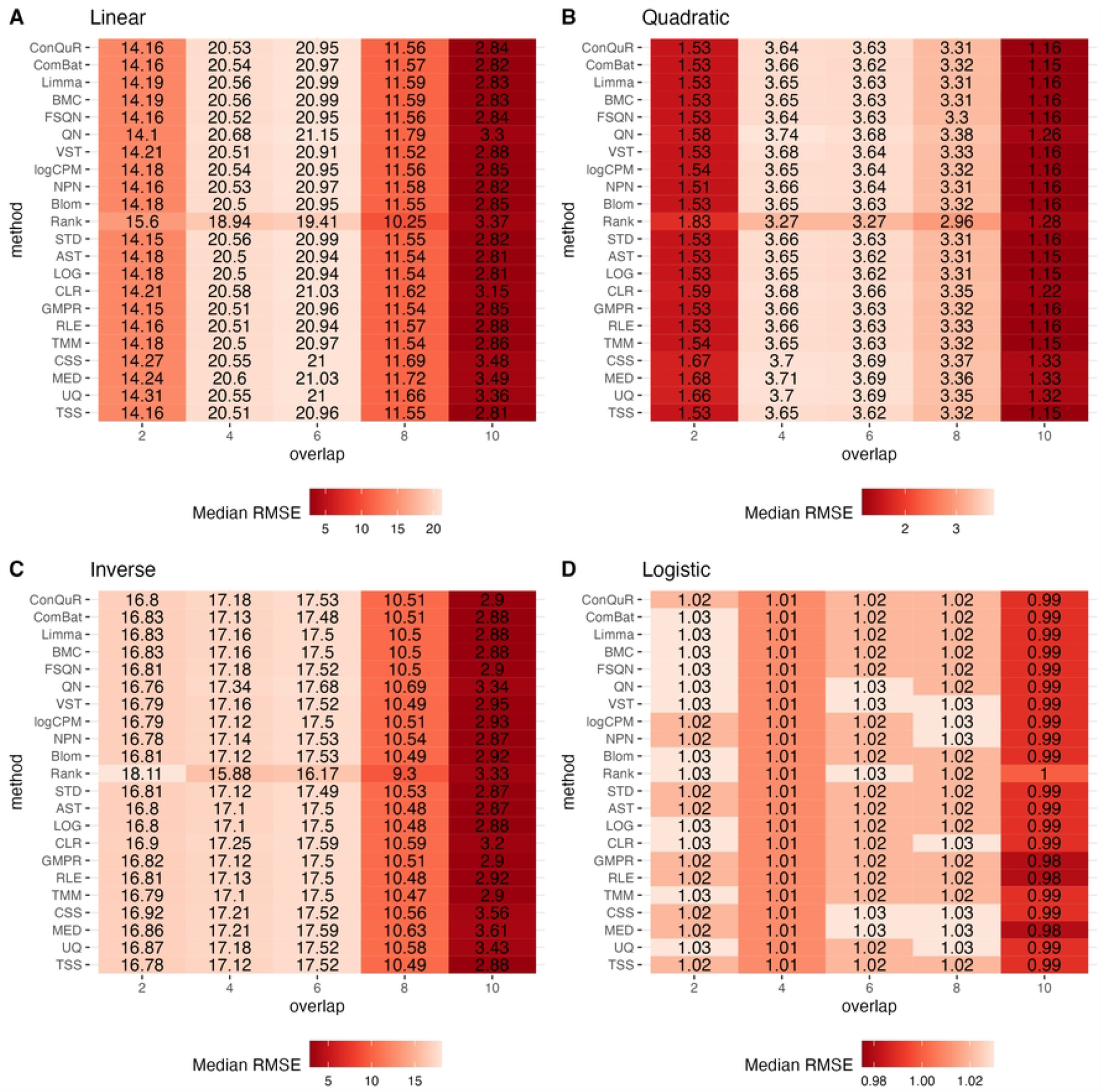
Heatmaps depicting median RMSE values obtained from abundance profiles normalized by different methods for predicting simulated quantitative phenotype in Scenario 3. The panels correspond to relationships between phenotype and phenotype-associated taxa, including A: linear, B: quadratic, C: inverse, and D: logistic. The columns represent different numbers of overlapping disease-associated taxa in the training and testing datasets. The rows represent different normalization methods.

In our simulations for predicting quantitative phenotypes, we consistently found that no normalization method exhibited consistent advantages over the others. However, given the frequent occurrence of batch effects and the satisfactory performance of batch correction methods in predicting datasets with such effects in both the training and testing sets, we highly recommend utilizing batch correction methods as an initial step prior to predicting quantitative phenotypes.

### Use QN and ComBat normalization carefully in quantitative phenotype prediction

In the following analysis, we assess the performance of different normalization methods using a set of 31 shotgun sequencing datasets obtained from healthy stool samples (Table 1). Each dataset is paired, with one assigned for model training and the other for validation purposes. For each method, we calculate the RMSE values based on the normalized abundance using a random forest model. To account for the randomness inherent in the prediction model, we repeat the predictions 10 times and report the median RMSE value for each study.

S3 Fig shows boxplots of the median RMSE obtained from predictions made using various models on specific test datasets. Within these specific test datasets, we performed Wilcoxon tests to evaluate the differences in means between different methods and the average mean. Our observations indicate that all methods encounter limitations due to biological and technical factors when predicting quantitative phenotypes, despite their best efforts. None of the methods exhibited significant reductions in the prediction’s RMSE, and no significant differences were observed among them. This aligns with the conclusions derived from our simulations. For instance, as shown in S3 FigC2, when KeohaneDM 2020 was used as the test dataset while others served as training sets, the RMSE values varied from 5.5 to 11.9 depending on the selected training data. The median RMSE values were approximately 8.2, without any significant differences observed among them.

To quantify the performance of normalization methods, we ranked all normalization methods based on the median RMSE values when the model was trained on the same dataset and validated on the same test dataset. Fig 5 shows the distributions of the ranks for each method across the 31 studies. A higher ranking (lower values in the box plot) indicates a better prediction performance. While all normalization methods had similar performance, batch correction methods exhibited slightly better results. It is worth mentioning that QN and ComBat, among the batch correction methods, displayed fluctuations that made them susceptible to extreme rankings compared to the other 22 normalization methods. Methods like CLR, LOG, and logCPM showed similar patterns. Therefore, caution should be exercised when employing these methods. Based on these findings, we suggest employing batch correction methods like FSQN, BMC, and Limma when making predictions for quantitative phenotypes.

**Fig 5.**
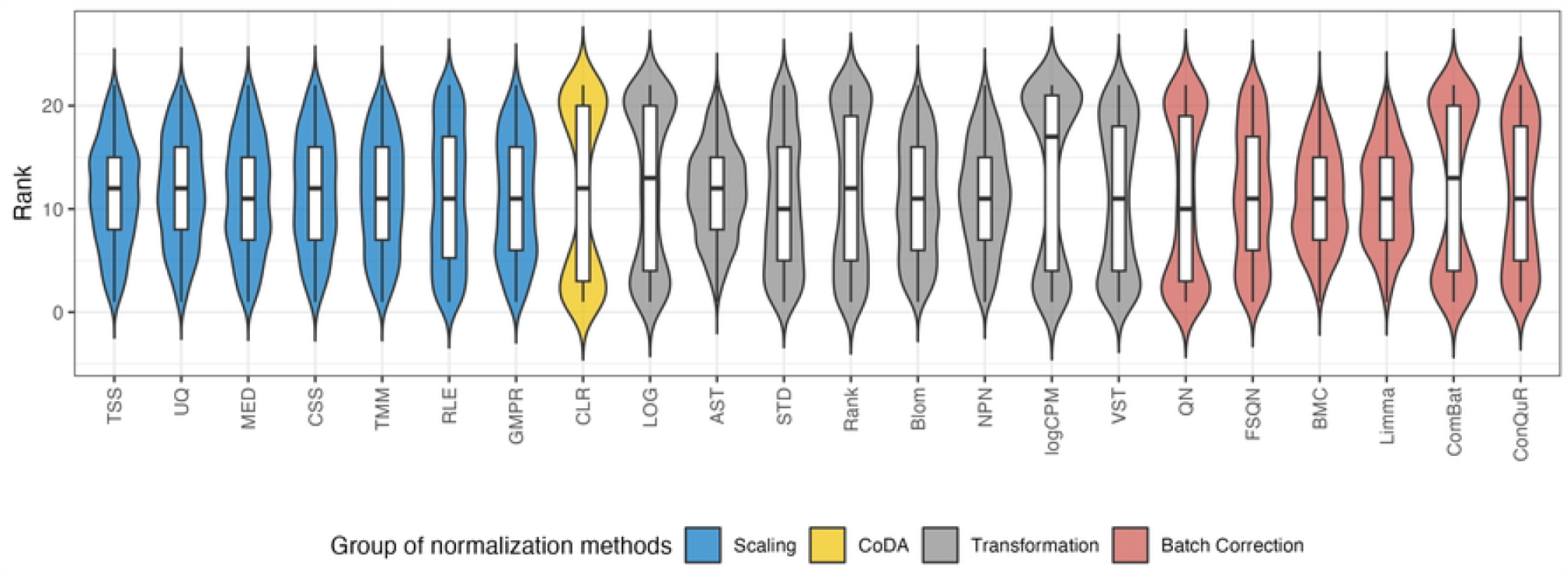
The distribution of the ranks for different normalization methods according to the median RMSE value under the same training and testing datasets.

## Discussion

The performance of normalization methods in metagenomic data analysis is a critical concern, particularly when predicting quantitative phenotypes. In our study, we investigated the performance of different normalization methods for predicting quantitative phenotypes using simulations and real datasets. Our investigation provides valuable insights into the performance and suitability of different normalization approaches for predicting quantitative phenotypes.

In our previous study of binary phenotype prediction [11], scaling methods such as TMM and RLE demonstrated relatively good performance, while transformation methods, including NPN and Blom, exhibited promising results in certain datasets. Furthermore, batch correction methods, such as BMC and Limma, consistently performed well across multiple datasets. The challenges encountered in predicting quantitative phenotypes were evident across all normalization methods, as none of them achieved a significant reduction in the RMSE of the predictions irrespective of the approach employed. These findings align with our simulations and underscore the complex nature of metagenomic data, which is prone to both biological and technical variations. Hence, it is reasonable to infer that the limitations are inherent to the data itself rather than being contingent on the choice of normalization method.

The absence of significant differences among the normalization methods is an important observation. Despite considering a wide range of relationships between phenotypes and taxa abundance profiles (linear, quadratic, inverse, and logistic), the variation in RMSE values remains consistently low across the methods. This resilience of normalization methods across various scenarios is a valuable finding as it enables researchers to choose methods based on other criteria without compromising predictive performance.

However, our analysis unveiled a modest advantage of batch correction methods over other normalization techniques. Specifically, we observed slightly improved results when employing these methods. Among the recommended methods for predicting quantitative phenotypes are FSQN, BMC, and Limma. Although they may not yield drastic performance enhancements, their slightly superior performance signifies potential robustness in tackling the inherent challenges of metagenomic data, particularly in the prediction of quantitative phenotypes.

It is crucial to acknowledge that certain normalization methods, namely QN and ComBat, exhibited fluctuations that heightened their susceptibility to extreme rankings. These fluctuations underscore the importance of exercising caution when selecting specific normalization techniques. Hence, researchers must carefully evaluate the suitability of a chosen method for their particular dataset and research question, taking into account the unique characteristics and potential fluctuations inherent in their data.

In conclusion, the performance of normalization methods in analyzing metagenomic data remains an active and ongoing area of research. Our study contributes to this field by conducting a comprehensive evaluation of various methods and providing valuable insights into their effectiveness in predicting quantitative phenotypes. From our findings, it appears that batch correction methods may be preferable. However, it is still crucial for researchers to continue exploring and developing novel techniques to further enhance the accuracy of predictions in the intricate realm of metagenomic data. Ultimately, the selection of a normalization method should be made judiciously, considering the specific characteristics of the dataset and the research objectives, as there is currently no universally applicable solution in this challenging domain.

## Supporting information

**S1 Fig. Boxplot illustrating the distribution of BMI values across different datasets**. he red dots indicate the average BMI for each dataset, and the gray dashed line represents the overall BMI mean. The significance of the difference between dataset BMIs and the overall BMI was determined using the Wilcoxon test, where ns signifies non-significance, ^*^ denotes a p-value *<* 0.05, ^**^ indicates p-value *<* 0.01, ^***^ suggests p-value *<* 0.001, and ^****^ represents p-value *<* 0.0001.

**S2 Fig. Boxplot illustrating the distribution of Shannon Index across different datasets**. The red dots indicate the average Shannon Index for each dataset, and the gray dashed line represents the overall average Shannon Index. The significance of the difference between dataset Shannon Indices and the overall Shannon Indices was determined using the Wilcoxon test, where ns signifies non-significance, ^*^ denotes a p-value *<* 0.05, ^**^ indicates p-value *<* 0.01, ^***^ suggests p-value *<* 0.001, and ^****^ represents p-value *<* 0.0001.

**S3 Fig. Predictive performance of different normalization methods using 31 healthy stool datasets from curatedMetagenomicData**. This figure comprises a collection of boxplots, each representing the predictive performance of different normalization methods in a specific test dataset. The x-axis enumerates various normalization techniques, while the y-axis indicates the median RMSE obtained from ten repeated predictions. The red dots signify the mean median RMSE for each method. The significance of differences between the method means and the overall mean was assessed by the Wilcoxon test, with “ns” for non-significant results.

## Acknowledgments Funding

This work was supported by the National Key R&D program of China [grant number 2018YFA0703900] and the National Science Foundation of China [grant number 11971264].

## Data availability

All the datasets used in this study are available in the R package curatedMetagenomicData (v3.8.0). All the codes used in the analysis can be found at https://github.com/wbb121/NormMethodsComp–QuantPred.

## Competing interests

The authors declare no competing interests.

## Author contributions

Y.L. designed and supervised the study. B.W. implemented the methods, conducted the computational analysis, and drafted the manuscripts. Y.L. modified and finalized the manuscripts. All authors read and approved the final version of the manuscript.

## Notes

### Competing Interest Statement

The authors have declared no competing interest.

